# In-depth proteomic profiling of the extracellular matrix of pancreatic ductal adenocarcinomas identifies signatures correlating with lymphocyte infiltration

**DOI:** 10.1101/2025.08.22.669728

**Authors:** James M. Considine, Dharma Pally, Sarah A. O’Brien, Jessica Potts, Di Feng, Jeanine Pignatelli, Abhishek S Kashyap, Nikita Sharma, Alexandra Naba

## Abstract

The extracellular matrix (ECM) is a complex assembly of proteins surrounding cells. It is a critical component of the tumor microenvironment that plays an active role in tumor progression and modulation of tumor response to treatment. Pancreatic ductal adenocarcinoma (PDAC) is a cancer type characterized by one of the worst prognoses, as it is often diagnosed at an advanced stage. It is also characterized by a very dense ECM, which hinders efficient drug delivery. In addition, PDACs are considered “cold” tumors as they fail to elicit a strong immune response, challenging the use of immunotherapy for PDAC cancer patients. Yet, the interplay between the ECM and immune cells within the PDAC tumor microenvironment remains poorly understood. Here, we employed ECM-focused proteomics to profile the ECM compositions of PDAC mouse models characterized by different levels of CD8^+^ T-cell infiltration. We found that CD8^lo^, or “cold” tumors, and CD8^hi^, or “hot” tumors, exhibited different ECM compositions. Interrogation of publicly available single-cell RNA-sequencing datasets of human PDACs further revealed that the ECM proteins distinguishing hot and cold PDACs are secreted by multiple stromal cell populations, including cancer-associated fibroblasts, stellate cells, and macrophages. Last, we found that the expression of a subset of the genes encoding ECM proteins characteristic of the CD8^lo^ phenotype correlated with CD8^+^ T-cell infiltration in human PDAC samples and patient survival. This study paves the way for the development of ECM-modulating interventions to enhance immune cell infiltration and responsiveness to immunotherapy.

**Significance:** We report the identification of ECM protein signatures correlating with the level of CD8^+^ lymphocyte infiltration in murine models of pancreatic ductal adenocarcinomas and human samples. This work paves the way for the development of ECM-modulating therapeutic strategies to enhance lymphocyte infiltration and, hence, the efficacy of immunotherapies.

## INTRODUCTION

Over the past two decades, significant advances in immunotherapy, a therapeutic modality that aims to stimulate a patient’s immune system - and in particular cytotoxic CD8^+^ lymphocytes (or CD8^+^ T cells) - to attack its tumor, have transformed the lives of countless cancer patients (1–5). These advances have been facilitated by an increased appreciation for the importance of the tumor microenvironment (TME), that is, the non-tumor-cell components of a tumor, in cancer progression and response to therapy (6–8).

A prominent component of the tumor microenvironment is the extracellular matrix (ECM). The ECM is a complex and dynamic assembly of over 150 proteins that constitutes a scaffold providing physical and mechanical support and signaling cues to cells (9). Most tumors are characterized by increased ECM content, or desmoplasia, and we know that changes in ECM composition or architecture affect all steps of cancer progression, from supporting tumor cell proliferation and survival, to promoting tumor angiogenesis and enhancing cell invasion and dissemination (10–12). We also now have evidence that the tumor ECM composition can be altered by anti-cancer treatments (13). Conversely, the ECM can influence the composition of the TME, in particular the immune microenvironment (14,15), and hence can modulate response to anti-cancer treatment (14,16,17). Thus, intervening on the ECM could increase the efficacy of anti-cancer therapies, including immunotherapy (18).

Pancreatic ductal adenocarcinoma (PDAC) is a cancer type characterized by one of the worst prognoses, as it is often diagnosed at an advanced stage. The American Cancer Society projects that this cancer will claim the lives of over 50,000 patients in the US in 2025 (19). PDAC is notoriously characterized by excessive ECM accumulation and high stromal cell content (including cancer-associated fibroblasts, stellate cells, and inflammatory cells). A recent study has shown that ECM content negatively correlates with PDAC patient survival (median survival of patients with ECM^hi^ tumors: 15.3 months vs. 22.9 months for patients with ECM^lo^ tumors (p=0.02) (20). A series of studies by Tian and collaborators using patient samples and mouse models of PDACs has identified subsets of ECM proteins correlating with PDAC progression and metastatic dissemination (21–23). It has thus been proposed that targeting components of the PDAC stroma by directly modulating the ECM or the cells contributing to the production of the ECM could offer long-awaited therapeutic benefits (24–27). Importantly, PDACs are often considered “cold”, *i.e.*, as failing to elicit a strong immune response, challenging the use of immunotherapy. Because of the demonstrated interplay between the tumor stroma, the ECM, and lymphocytes, and the successes of immunotherapies, it thus becomes appealing to attempt to modulate the PDAC stroma and the ECM to convert a cold TME into a hot one and render these tumors responsive to immunotherapy (28–30). However, to do so, we first must identify relevant ECM targets.

Here, we used proteomics to profile the ECM composition of *KRAS*-driven and translationally relevant murine models of CD8^hi^ (“hot”) and CD8^lo^ (“cold”) PDACs. Using highly stringent identification criteria, we report the identification of a 16-ECM-protein signature characteristic of a cold phenotype and an 8-ECM-protein signature correlating with a hot tumor phenotype. We validated these findings using immunohistochemistry, which provides additional insights into the distribution pattern of ECM proteins within the PDAC TME. Leveraging publicly available single-cell RNA sequencing datasets generated on human PDAC samples, we further demonstrate that the genes encoding signature ECM proteins are expressed by multiple cell populations of the PDAC TME and that their expression levels correlated with CD8^+^ T cell infiltration. Last, we show that ECM gene expression correlates with PDAC patient survival. Our work paves the way for the development of matritherapies (31) that will aim to modulate the composition of the ECM to attempt to increase immune cell infiltration and, hence, increase immunotherapy efficacy. Importantly, with an in-depth characterization of the matrisome of hot PDACs, we can also begin to design approaches using ECM proteins for the targeted delivery and anchoring of therapeutic payloads for enhanced efficacy and reduced side effects.

## MATERIALS AND METHODS

### Cell lines and animal studies

All animal studies were performed in accordance with Boehringer Ingelheim’s Institute for Animal Care and Use Committee (IACUC protocol number 19-636-E). Female C57BL/6 mice were purchased from Jackson Laboratories and used at 6-8 weeks of age. KPCY TC3 (6419c5), KPCY TC4 (6422c1), KPCY TH2 (2838c3), and KPCY TH3 (6499c4) cell lines are derived from a previously established autochthonous mouse model of pancreatic ductal adenocarcinoma (*Kras^LSL-G12D/+^*;*Trp53^LSL-R172H/+^*;*Pdx1-Cre*; *Rosa26^YFP/YFP^*, referred to as “KPCY PDAC”) (32,33) and were acquired from Dr. Stanger at the University of Pennsylvania. For mouse tumor studies, 3.0×10^5^ cells were injected subcutaneously in the right flank. Tumors were measured three times per week, and tumor volume was calculated using the following formula: [(length x width^2^ x π)/6]. Tumors were harvested at 500 mm^3^ to compare immune infiltration by immunostaining and harvested at >100 mm^3^ and snap frozen in liquid nitrogen for subsequent proteomic analyses.

### Histological staining

Mouse tumors were fixed in formalin and embedded in paraffin according to standard procedures. 5-μm sections were mounted on positively charged slides, dried, and baked at 60°C for an hour. Sections were deparaffinized using xylene and rehydrated by incubation in solutions of descending ethanol concentrations and then in tap water. Adjacent sections were stained with the following stains:

### Masson’s trichrome staining

Deparaffinized sections were incubated in Bouin’s fixative (StatLab, #FXBOULT) for 1 hour at 56°C, rinsed with tap water, stained with Biebrich Scarlet solution (Electron Microscopy Sciences, #26033-25) for 8 min. at room temperature, washed in distilled water, incubated in phosphotungstic/phosphomolybdic acid (Electron Microscopy Sciences, #2636705) for 15 min. and finally, stained with Aniline Blue solution (Electron Microscopy Sciences, #2636706) for 10 min.

#### Picrosirius red staining

Deparaffinized sections were incubated in Bouin’s fixative (Mercedes Scientific, #EKI 22771L) for 1 h at 60°C, rinsed with tap water, stained with 0.1% Fast Green counterstain for 15 min. at room temperature, incubated with 1% acetic acid for 4 min., and finally, stained with Picrosirius red solution (Mercedes Scientific, #SRS999) for 30 min. Slides were briefly rinsed in distilled water, dehydrated, and mounted with Micromount (Leica Microsystems, #3801730).

#### Immunohistochemistry

The following antibodies were used: anti-α smooth muscle actin (αSMA), anti-CD8, anti-collagen VIII (Col8a1), anti-fibulin 5, anti-MMP9, anti-periostin, and anti-TGFβ induced. Catalog numbers, concentrations, and epitope retrieval conditions used are provided in **Supplementary Table S2**. Staining was performed with the BOND Polymer Refine Detection Kit (Leica, #DS9800) on the BOND RX automated stainer (Leica Biosystems) according to a standard preset protocol. After deparaffinization, sections were subjected to heat-induced epitope retrieval. Endogenous peroxidase activity and non-specific binding sites were blocked by sequentially treating samples with peroxidase block (BOND Polymer Refine Detection Kit) and protein block (Background Sniper, Biocare Medical, #BS966) for 15 min. at room temperature. Sections were then incubated with the primary antibody for 30 min. After several washes, signal detection was performed with a horseradish peroxidase (HRP)-coupled anti-rabbit secondary antibody and 3, 3’-diaminobenzidine (DAB; BOND Polymer Refine Kit) by incubating the sections for 15 min. and 10 min. at room temperature, respectively. Tumor sections were counterstained with hematoxylin. All slides were dehydrated in an Autostainer XL and mounted with Micromount (Leica Biosystems).

### Histological analysis using HALO

All slides were scanned at a resolution of 40x on a Leica Aperio AT2 whole slide scanner. Regions of interest were delineated for each tumor section and manually annotated using Aperio ImageScope (v.12.4.6.5003) to exclude artifacts such as tissue folds. Images and annotations were imported to HALO (Indica Labs) and analyzed as described below.

#### Determination of CD8^+^ T-cell density

Hematoxylin and DAB stains were manually assigned in the algorithm, and thresholds were adjusted to detect and differentiate nuclei and CD8^+^ T cells. After obtaining the number of CD8^+^ T cells for each tumor section, we calculated the density of CD8^+^ T cells by dividing the number of CD8^+^ T cells by the tumor area in mm^2^.

#### Quantification of SMA-positive (SMA^+^) staining

First, tumor areas were identified by training a random forest classifier algorithm to separate tumor tissue from any surrounding stroma, necrosis, tissue folds, and background whitespace areas. Once tumor areas were identified, the analysis proceeded to assess SMA^+^ staining by means of thresholding. The SMA^+^ staining area was determined using the HALO area quantification algorithm (v2.3.1) by first defining the settings for the hematoxylin counterstain, followed by setting thresholds to detect the aSMA stain positivity of weak, moderate, and strong signal. The results were expressed as total areas of tissue analyzed, areas of positive staining, and percentage positive areas, which are normalized measures considering the total tissue areas analyzed for each slide.

#### Determination of tumor collagen content

The area quantification HALO area quantification algorithm was used to quantify the percentage of collagen area in tumor sections stained with either Masson’s trichrome or picrosirius red. Color channels were manually assigned by selecting single-stained regions. A third channel was added to recognize and remove artifactual stains (black or brown) and was labeled “exclusion stain”. Thresholds were set manually for the two primary colors in each image. Two phenotypes were defined: a tissue-positive region where collagen staining is negative, but counterstain is positive, and a collagen-positive region where collagen staining is positive (blue when using Masson’s trichrome and green when using picrosirius red). Collagen content is calculated by dividing the collagen-positive area by the total tumor area (*i.e.*, tissue-positive and collagen-positive regions). Statistical analysis was performed using the Welch’s *t*-test, assuming unequal variance for all conditions.

### Sample preparation for proteomic analysis

#### Tissue decellularization

40 to 120 mg of tissue were homogenized using an OMNI Bead Ruptor using the 8 m/s setting with three 10-sec. cycles and a 60-sec. break between each cycle. The bead mix used for tissue homogenization contained four 2.8-mm ceramic beads (OMNI International #19628) and four 2.4-mm metal beads (OMNI International #19-610). ECM enrichment was achieved through the sequential extraction of intracellular proteins in order of decreasing protein solubility, using a subcellular protein fractionation kit for tissues following the manufacturer’s instructions (Thermo Scientific, #87790) as previously described (34). The efficiency of the sequential extraction of intracellular components and concomitant ECM-protein enrichment was monitored by western blot analysis using the following antibodies: anti-collagen I (Sigma, #AB765P, used at a 0.5 µg/mL), anti-histone H4 (Sigma, #05-858, antibody used at 1/30,000 dilution), anti-GAPDH (Sigma, #MAB374, used at 2 µg/mL), and anti-integrin beta-1 serum kindly gifted by Dr. Richard O. Hynes (used at 1/1000 dilution).

We analyzed five independent tumors for each tumor model for a total of ten tumors with low CD8^+^ T-cell infiltration (CD8^lo^), also referred to as “cold” tumors” (TC3, n = 5 and TC4, n = 5), and ten tumors with high CD8^+^ T-cell infiltration (CD8^hi^), also referred to as “hot” tumors (TH2, n = 5 and TH3, n = 5).

#### Protein digestion for mass spectrometry analysis

ECM-enriched protein samples were subsequently solubilized and digested into peptides following an established protocol (35–37). In brief, proteins were solubilized in an 8 M urea solution prepared in 100 mM NH_4_HCO_3_, and protein disulfide bonds were reduced using 10 mM dithiothreitol (Thermo Scientific, #A39255). Reduced disulfide bonds were then alkylated with 25 mM iodoacetamide (Thermo Scientific, #A39271) for 30 min. in the dark, at room temperature. The urea concentration was brought to 2 M, and proteins were then deglycosylated with PNGaseF (New England Biolabs, #P0704L) for 2 hours at 37°C and digested sequentially, first with Lys-C (Thermo Scientific, #90307) for 2 hours at 37°C, and then with trypsin (Thermo Scientific, #90058), overnight at 37°C. A fresh aliquot of trypsin was added the following day and samples were incubated for an additional 2 hours at 37°C. All incubations were performed under mild agitation. Samples were acidified with 50% trifluoroacetic acid (TFA) and desalted using Pierce Peptide Desalting Spin Columns, as previously described (38). Peptides were lyophilized and then reconstituted in a solution containing 95% HPLC-grade water, 5% acetonitrile, and 0.1% formic acid, and the concentration of the peptide solution was measured using the Pierce Quantitative Colorimetric Peptide Assay kit (#23275).

#### Peptide fractionation

40 μg of each sample were fractionated by high-pH reversed-phase chromatography using Shimadzu UFLC (Kyoto, Japan) and Waters (Milford, US) xbridge column (C18 4.6 x 150mm, 3.5 μm). Buffer A consisted of 10 mM ammonium formate and buffer B consisted of 10 mM ammonium formate with 90% acetonitrile; both buffers were adjusted to pH 10 with ammonium hydroxide. The gradient starts with 1% B at 1 mL/min for 3 min., then 1% B to 25% B in 60 min., 65% B in 10 min., and ramped to 85% B in 5 min. The gradient was held at 85% B for 10 min. before being ramped back to 1% B. 80 fractions were collected at 1mL/min. flow rate and combined into 8 pools as follows: first, the last 8 fractions (fractions 73-80) were pooled with fractions 65-72, with fraction 80 being combined with 72, 79 with 71, 78 with 70, etc. The volume of these pooled fractions was brought down using a SpeedVac to 1 mL each and pooled with the next set of fractions, 72 being pooled with 64, 71 with fraction 63, and 70 with 62, etc. Sample volume was reduced to 1 mL and samples were then pooled with the next set of 8 fractions, 64 was pooled with fraction 56, 63 with 55, and 62 with 54, etc. This process was repeated until the concatenation of 80 fractions into 8 pools, each of 10 fractions, was achieved. Concatenation was followed by desalting using Nestgroup C18 tips (Southborough, MA). Fractionated peptides were dried and redissolved in 25μL 0.1% FA, and each pool was analyzed using LC−MS/MS. All reagents were LC/MS grade and purchased from Sigma-Aldrich. All solvents were Optima LC/MS-grade and purchased from Fisher Chemical.

#### LC-MS/MS data acquisition

Approximately 5 μg of pooled fractionated peptides were analyzed using a Q Exactive HF mass spectrometer coupled with an UltiMate 3000 RSLC nanosystem with a Nanospray Flex Ion Source (Thermo Fisher Scientific). The samples were loaded into a Waters nanoEase M/Z C18 (100Å, 5um, 180um x 20mm) trap column and then a 75μm x 150mm Waters BEH C18 (130A, 1.7um, 75um x 15cm) column and separated at a flow rate of 300 nL/min. Solvent A was 0.1% FA in water and solvent B was 0.1% FA, 80% ACN in water. The solvent gradient of LC was 5% B in 0-3 min., 10% B at 3.2 min., 40% B at 55 min., 95% B at 60 min., wash 95% for 5 min., followed by 5% B equilibration until 75 min. Full MS scans were acquired in the Q-Exactive HF mass spectrometer over the 350-1400 m/z range with a resolution of 60,000. The AGC target value was 1.00E+06 for the full scan. The 15 most intense peaks with charge states 2, 3, 4, 5 were fragmented in the HCD collision cell with a normalized collision energy of 30%; these peaks were then excluded for 30s within a mass window of 1.2 m/z. A tandem mass spectrum was acquired in the mass analyzer with a resolution of 15,000. The AGC target value was 5.00E+04. The ion selection threshold was 1.00E+04 counts, and the maximum allowed ion injection time was 30 msec. for full scans and 50 msec. for fragment ion scans.

#### Database searching

Spectra were searched against the UniProt mouse database (0221207_Uniprot mouse containing 17,138 entries) using MaxQuant with the following parameters: parent mass tolerance of 20 ppm, fragment ion mass tolerance of 20ppm, and assuming the digestion enzyme stricttrypsin and allowing a maximum of 2 missed cleavage sites. Carbamidomethyl (C) of cysteine was specified as a fixed modification. Gln->pyro-Glu of the N-terminus, deamidation of asparagine and glutamine, and oxidation of methionine, proline, and lysine were specified as variable modifications, the latter two being characteristic post-translational modifications of ECM proteins, in particular collagens and collagen-domain-containing proteins, as we previously reported (39). MaxQuant LFQ was used for label-free quantification using the default setting.

#### Criteria for protein identification and data analysis

Scaffold (version Scaffold_5.2.1, Proteome Software Inc., Portland, OR) was used to validate MS/MS-based peptide and protein identifications. We used stringent criteria to accept peptide and protein identification: peptide identifications were accepted if they could be established to achieve an FDR of less than 1.0% by the Percolator posterior error probability calculation (40); protein identifications were accepted if they could be established with an FDR of less than 1.0% and at least 2 identified peptides. Protein probabilities were assigned by the Protein Prophet algorithm (41). Proteins that contained similar peptides and could not be differentiated based on MS/MS analysis alone were grouped to satisfy the principles of parsimony. Proteins sharing significant peptide evidence were grouped into clusters.

Mass spectrometry output was further annotated to identify ECM and non-ECM components (35,42). Specifically, matrisome components were classified as core-matrisome or matrisome-associated components and further categorized into groups based on structural or functional features: ECM glycoproteins, collagens, or proteoglycans for core matrisome components; and ECM-affiliated proteins, ECM regulators, or secreted factors for matrisome-associated components. Total precursor ion intensities were used to estimate protein abundance and conduct label-free inter-group quantification. Unpaired Student’s t-test was applied to determine the statistical significance of changes in protein abundance between CD8^hi^ and CD8^lo^.

### Single-cell RNA sequencing

A published single-cell RNA sequencing dataset of human pancreatic ductal adenocarcinoma (43) was retrieved from the China National Center for Bioinformation (https://ngdc.cncb.ac.cn/gsa/) with the accession number GSA: CRA001160. Scanpy 1.9 was used to analyze and visualize the scRNA-seq datasets in this study, keeping the t-SNE coordinate provided in the original publication.

### ssGSEA and Tumor IMmune Estimation Resource (TIMER) analyses

To examine the relationship between gene expression and immune cell infiltration, we used single-sample gene set analysis (ssGSEA) (44) and the TIMER algorithm via the TIMER2.0 web server (http://timer.cistrome.org/) (45). The Gene module in TIMER2.0 was used to calculate the association score of each gene with CD8^+^ T cells. A purity adjustment was used to account for the influence of tumor purity on the relationship between ECM gene expression and immune cell infiltration levels in the TCGA PAAD (PDAC) RNA-Seq dataset. Associations were considered significant if the p-value was less than 0.05.

### Survival analysis

We used the open-source TCGA2STAT R package v 1.2 to obtain level 3 TCGA gene expression data (summarized as RSEM value) from the Xena TOIL TCGA dataset and corresponding clinical annotations. TCGA2STAT was used to prepare data into a format ready for statistical analysis using TCGA survival. First, each gene of interest was examined for its effect on survival by separating patients into high/low expression subgroups. Maximally Selected Rank Statistics (R package maxstat) was used to estimate the best gene expression cutoff that separates high/low expression subgroups with differential survival. We also utilized single-sample gene set enrichment analysis (ssGSEA) to categorize pancreatic ductal adenocarcinoma patients from the TCGA data (https://portal.gdc.cancer.gov/) based on their level of expression of the matrisome genes of the CD8^hi^ or CD8^lo^ signatures. Gene set score was examined for its effect on survival by separating patients into high/low score subgroups based on the median value. We applied the Cox proportional hazards model to determine the hazard ratio (HR) and p-value between the high gene-signature expression group and low gene-signature expression group. The R packages used in this analysis included survminer and survival.

### Data visualization

All graphs were plotted using PRISM (GraphPad). Proportional Venn diagrams were generated using BioVenn: https://www.biovenn.nl/ (46). The intersection of multiple datasets was visualized using UpSetR and the online application: https://gehlenborglab.shinyapps.io/upsetr/ (47). Network and pathway analysis were performed using STRING v12: https://string-db.org/ (48). Panels in Figure 1A and 1E were generated using BioRender: https://www.biorender.com/.

**Figure 1.**
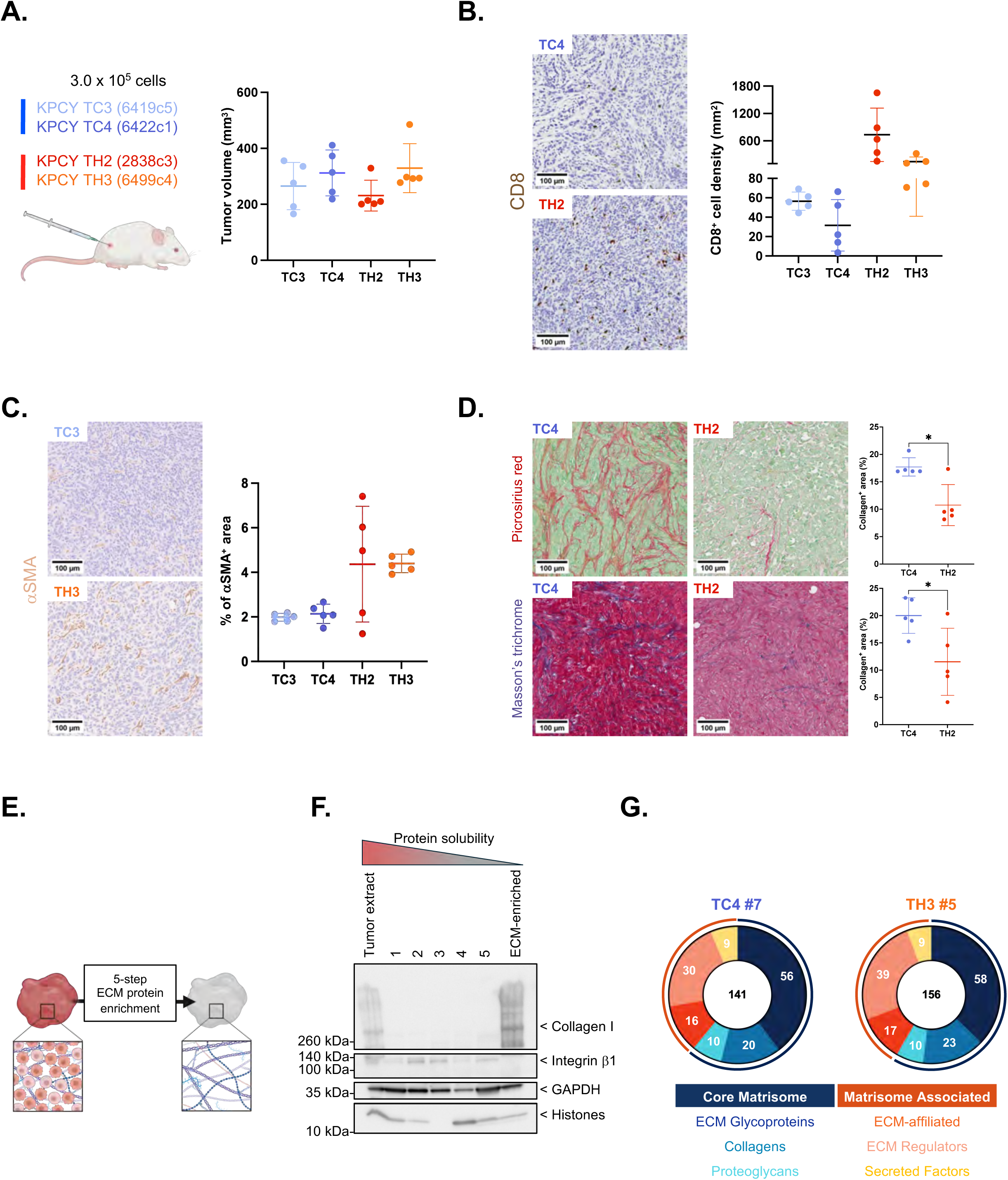
Experimental design and model system. **A.** Dot plot represents the tumor volumes of each sample (n=5/group) at experimental endpoint. The average tumor volume for each tumor group is indicated by a bar and the error bars represent the standard deviation (SD). Schematic was generated using BioRender. **B.** Two sections of each tumor sample (n=5/group) were stained using an anti-CD8 antibody. Representative images of CD8 staining of CD8^lo^ (TC4) and CD8^hi^ (TH2) tumors are shown (scale bar = 100μm). Dot plot represents the density of CD8^+^ T cells per tumor area (average of two sections from each tumor); data for each tumor group is indicated by a bar and the error bars represent the standard deviation (SD). **C.** Two sections of each tumor sample (n=5/group) were stained using an anti-αSMA antibody. Representative images of αSMA staining of CD8^lo^ (TC3) and CD8^hi^ (TH3) tumors are shown (scale bar = 100μm). Dot plot represents the proportion of SMA-positive tumor area. Data for each tumor group is indicated by a bar and the error bars represent the standard deviation (SD). **D.** Two sections of each tumor sample (n = 5/group) were stained using picrosirius red (PSR; top panels) or Masson’s trichrome to assess fibrillar collagen content. Representative images of picrosirius red staining (top panels) and Masson’s trichrome staining (bottom panels) of CD8^lo^ (TC4) and CD8^hi^ (TH2) tumors are shown (scale bar = 100μm). Dot plot represents the proportion of picrosirius-red-positive and Masson’s-trichrome-positive area (*i.e.*, “collagen-positive” area). The average positivity for each tumor group is indicated by a bar and the error bars represent the standard deviation (SD). Statistical analysis was performed using the Welch’s *t*-test, assuming unequal variance, and statistical significance is depicted as follows: * p <0.05. **E.** Schematic depiction of the experimental workflow leading to the enrichment of ECM proteins from tumors. Schematic was generated using BioRender. **F.** Representative immunoblots illustrate the sequential extraction of soluble non-ECM proteins (integrin β1, GAPDH, and histones) and enrichment of ECM proteins (collagen I) in the “ECM-enriched” protein fraction. **G.** Donut charts illustrate the distribution of matrisome proteins identified by mass spectrometry across matrisome categories in representative samples of CD8^lo^ (sample TC4#7) and CD8^hi^ (sample TH3#5) tumors. Core matrisome proteins include ECM glycoproteins (dark blue), collagens (light blue), and proteoglycans (teal); matrisome-associated proteins include ECM-affiliated proteins (dark orange), ECM regulators, including ECM-remodeling enzymes and their inhibitors (light orange), and secreted factors, including ECM-bound growth factors and cytokines (yellow).

### Data availability

Raw mass spectrometry data and the accompanying metadata file in the Sample and Data Relationship Format (SDRF) (49) have been deposited to the ProteomeXchange Consortium (50) via the PRIDE partner repository (51) with the dataset identifier PXD060932. The raw data will be made publicly available upon acceptance of the manuscript.

## RESULTS

### Characterization of the stromal composition of CD8^lo^ and CD8^hi^ subcutaneous KPCY PDAC

To study the possible correlation between immune-cell infiltration and ECM composition, we used a panel of previously established and characterized pancreatic ductal adenocarcinoma (PDAC) cell lines derived from a genetically-engineered mouse model expressing Kras^LSL-G12D/+^;Trp53^LSL-R172H/+^;Pdx1-Cre; Rosa26^YFP/YFP^, (referred to as KPCY PDAC) (32). We selected to work with two cell lines, KPCY TC3 (6419c5) and KPCY TC4 (6422c1), that when injected in mice form tumors with low CD8^+^ lymphocyte infiltration (further termed, CD8^lo^ or “cold” tumors) and with two cell lines, KPCY TH2 (2838c3) and KPCY TH3 (6499c4), that when injected in mice form tumors with higher CD8^+^ lymphocyte infiltration (further termed, CD8^hi^ or “hot” tumors) (**Figure 1A**). Importantly, we confirmed that the cell lines retained their phenotype when injected subcutaneously (**Figure 1B**). We also observed broader microenvironmental changes with CD8^lo^ tumors characterized by a smaller number of αSMA^+^ cancer-associated fibroblasts (CAFs) (**Figure 1C**). Last, we found that CD8^lo^ tumors have a statistically significantly higher ECM content as compared to CD8^hi^ tumors based on two histological stains specific for fibrillar collagens, picrosirius red (**Figure 1D, top panel**) and Masson’s trichrome (**Figure 1D, bottom panel**). The observation that CD8^lo^ tumors have higher fibrillar collagen content and yet less SMA^+^cells, often viewed as the main producer of collagens in the TME, suggests that either the CAFs in CD8^lo^ tumors produce more fibrillar collagens, that other cell populations within the CD8^lo^ TME produce less fibrillar collagens, or perhaps that the collagen ECM of the CD8^lo^ TME is more stable and less remodeled.

### Proteomics identifies signatures correlating with CD8 lymphocyte infiltration

To determine the protein composition of the ECM of CD8^lo^ and CD8^hi^ tumors, we first decellularized the tumors to achieve an enrichment of ECM proteins (**Figure 1E**). Using immunoblotting, we confirmed the extraction of intracellular proteins over the course of the five-step decellularization process and the resulting enrichment of ECM components (**Figure 1F**). ECM-enriched samples were subjected to proteolytic digestion, and the resulting peptides were fractionated using basic reversed-phase chromatography to achieve deep matrisome coverage and submitted to proteomic analysis using liquid chromatography coupled to tandem mass spectrometry (LC-MS/MS). We then used highly stringent criteria to accept peptide identification (<0.1% FDR) and protein identification (<1% FDR; detection with at least 2 unique peptides; **Supplementary Table S1A**). On average, we identified ∼150 distinct matrisome proteins in each tumor sample (**Supplementary Table S1B**) and found a consistent distribution of these proteins across the different categories of matrisome proteins, with ∼60% of the proteins belonging to core, structural, matrisome categories, including ECM glycoproteins, collagens, and proteoglycans, and ∼40% classified as matrisome-associated proteins (*i.e.*, ECM-affiliated proteins, ECM regulators including remodeling enzymes, and secreted factors including ECM-bound growth factors or cytokines) (**Figure 1G**). The top five most abundant ECM proteins were the same between the two tumor groups: collagen I (Col1a1 and Col1a2), collagen III (Col3a1), fibronectin (Fn1), and tenascin C (Tnc) (**Supplementary Table S1B**). The fact that this subset of proteins is almost two orders of magnitude more abundant than any of the other matrisome proteins detected likely explains why we were unable to identify differences between hot and cold tumors, as we have observed by immunohistochemistry.

We then defined the matrisome profile of each tumor model (TC3, TC4, TH2, and TH3) as the ensemble of proteins detected in at least three of the five independent biological replicates of each tumor model (**Supplementary Table S1C**). Intra-group comparison revealed a remarkable overlap between each biological replicate within each tumor model (**Supplementary Figure 1**), validating the robustness of our experimental pipeline. Similarly, inter-group comparison revealed a large overlap (130 proteins) between the matrisome profiles of the two CD8^lo^ tumor models (*i.e.*, TC3 vs TC4; **Figure 2A**). A similar proportion was observed when comparing the matrisome profiles of the two CD8^hi^ tumor models (*i.e.*, TH2 vs TH3; **Figure 2C**). Of note, the matrisome profiles of CD8^lo^ and CD8^hi^ tumors presented a similar distribution across matrisome categories, with the largest number of proteins belonging to the ECM glycoprotein, collagen, and ECM regulator categories (**Figures 2B and 2D, respectively**).

**Figure 2.**
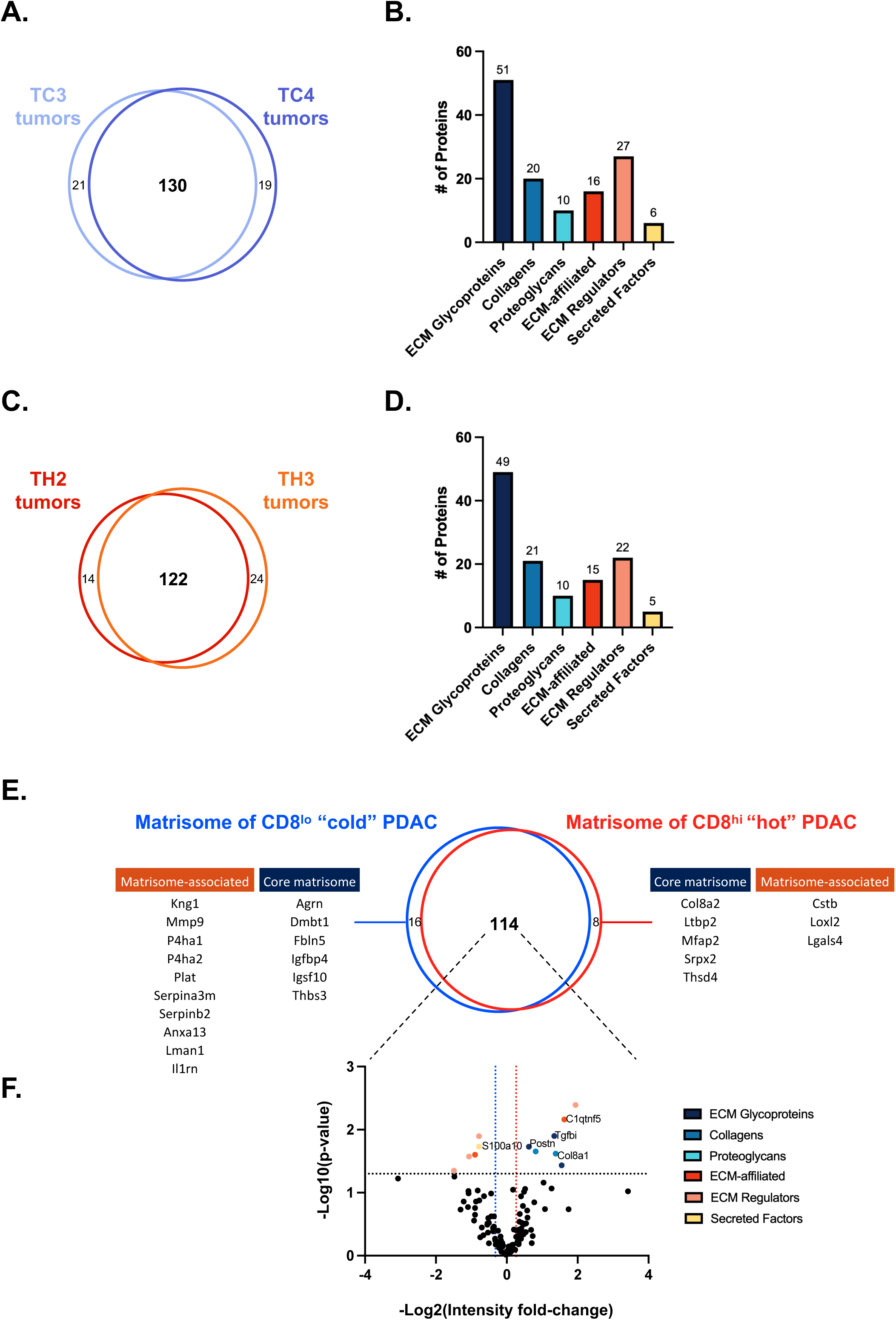
Proteomic definition of the matrisome of CD8^lo^ and CD8^hi^ KPCY tumors. **A.** Venn diagram represents the overlap between the matrisome proteins detected in at least three of the five independent biological replicates of the TC3 (light blue) and TC4 (dark blue) CD8^lo^ KPCY tumors. The 130 matrisome proteins detected in both tumor groups based on these stringent inclusion criteria constitute the matrisome of CD8^lo^ tumors. **B.** Bar graph represents the distribution of the 130 matrisome proteins detected in both tumor groups across the different matrisome categories. **C.** Venn diagram represents the overlap between the matrisome proteins detected in at least three of the five biological replicates of the TH2 (red) and TH3 (orange) CD8^hi^ KPCY tumors. The 122 matrisome proteins detected in both tumor groups based on these stringent inclusion criteria constitute the matrisome of CD8^hi^ tumors. **D.** Bar graph represents the distribution of the 122 matrisome proteins detected in both tumor groups across the different matrisome categories. **E.** Venn diagram represents the overlap between the matrisome of CD8^lo^ (blue) and CD8^hi^ (red) tumors defined above (see Figure 2A and 2C, respectively). **F.** Volcano plot represents the change in normalized total precursor ion intensity (*i.e.*, protein abundance) of the 114 matrisome proteins detected in both CD8^lo^ and CD8^hi^ proteins. Proteins detected in statistically differential abundance (p < 0.05 and ± 20% fold change) are depicted in the upper left and right quadrant and color-coded based on the matrisome category they belong to.

Last, we defined the matrisome profile of CD8^lo^ tumors and CD8^hi^ tumors as the ensemble of proteins detected in at least three of the five independent biological replicates and in both tumor models for a given tumor type. The comparison of these profiles identified 114 matrisome proteins detected in both tumor types (**Figure 2E**). Importantly, we identified a set of 16 matrisome proteins characteristic of CD8^lo^ tumors (**Figure 2E, left panel**). This set included six core matrisome components: agrin (Agrn), Dmbt1, fibulin 5 (Fbln5), insulin growth factor binding protein 4 (Igfbp4), Igsf10, and thrombospondin 3 (Thbs3) and ten matrisome-associated components including the matrix metalloproteinase 9 (Mmp9), an enzyme known to degrade collagens, and, interestingly, two prolyl hydroxylases, P4ha1 and P4ha2. Prolyl hydroxylases are the enzymes responsible for the addition of hydroxyl groups on prolines in glycine-X-Y amino acid motifs, prominent in collagens, a post-translational modification contributing to the stability of the collagen triple helix and hence of the entire collagen meshwork within the ECM (52). The observation that both enzymes are detected in higher abundance in the ECM of CD8^lo^ tumors may contribute to explaining our early observation that these tumors have relatively higher fibrillar collagen content than CD8^hi^ tumors. It also suggests that collagen synthesis and stability may be higher than MMP9-mediated collagen degradation in cold tumors.

We also identified a set of eight matrisome proteins characteristic of CD8^hi^ tumors (**Figure 2E, right panel**). This set comprised five core matrisome proteins and three matrisome-associated proteins, including the ECM crosslinking enzyme lysyl oxidase-like 2 (Loxl2), previously shown to associate with PDAC progression and the focus of active research and development efforts (53–56), demonstrating the rigor of our approach. It has previously been proposed that crosslinked, stiffer ECM hinders immune cell infiltration and function. Yet, we found that Loxl2 is present in higher abundance in CD8^hi^ tumors. These two somewhat opposite observations can be reconciled by our earlier observation that the fibrillar collagen content, substrate of Loxl2-mediated crosslinking, is lower in CD8^hi^ tumors, and so rather than crosslinking, the enzyme-to-substrate ratio is the factor permitting or limiting factor to lymphocyte infiltration.

Quantitative analysis of the proteins detected in the matrisome of both tumor types using normalized total precursor ion intensity to infer protein abundance identified an additional set of proteins present in statistically significant higher abundance in CD8^lo^ (**Figure 2F, top left quadrant**) such as S100a10, or CD8^hi^ tumors (**Figure 2F, top right quadrant**), such as periostin (Postn), the α1 chain of collagen VIII (Col8a1), or the TGFβ induced protein (Tgfbi). Altogether, these results define a 21-matrisome-protein signature characteristic of CD8^lo^ tumors and a 15-matrisome-protein signature characteristic of CD8^hi^ tumors.

### Immunostaining reveals the pattern of organization of the ECM in CD8^lo^ and CD8^hi^ PDAC tumors

Next, we selected a subset of proteins for which validated antibodies were available to confirm the proteomic results. In agreement with our proteomic data, fibulin 5 (**Figure 3A**) and Mmp9 (**Figure 3B**) presented a distinctive extracellular distribution in the microenvironment of CD8^lo^ tumors but were not detected in CD8^hi^ tumors. Conversely, collagen VIII presented a clear extracellular distribution in the microenvironment of CD8^hi^ tumors but was not detected in CD8^lo^ tumors. (**Figure 3C**). Interestingly, while we detected positive staining for the protein TGFβ induced in both CD8^lo^ and CD8^hi^ tumors (**Figure 3D**), the pattern was clearly intracellular in CD8^lo^ tumors and extracellular in CD8^hi^ tumors. We were further able to confirm this phenotype by assessing the presence of TGFβ induced by immunoblot and found that while TGFβ induced was detected in protein samples extracted from CD8^lo^ tumors, it was enriched in a protein fraction of high solubility, so likely intracellular (**Figure 3D, lower left panel**). In contrast, in CD8^hi^ tumors, TGFβ induced was retained in the ECM-enriched protein fraction characterized by high insolubility (**Figure 3D, lower right panel)**. These examples further validate the robustness of our discovery pipeline to identify ECM proteins of the tumor microenvironment.

**Figure 3.**
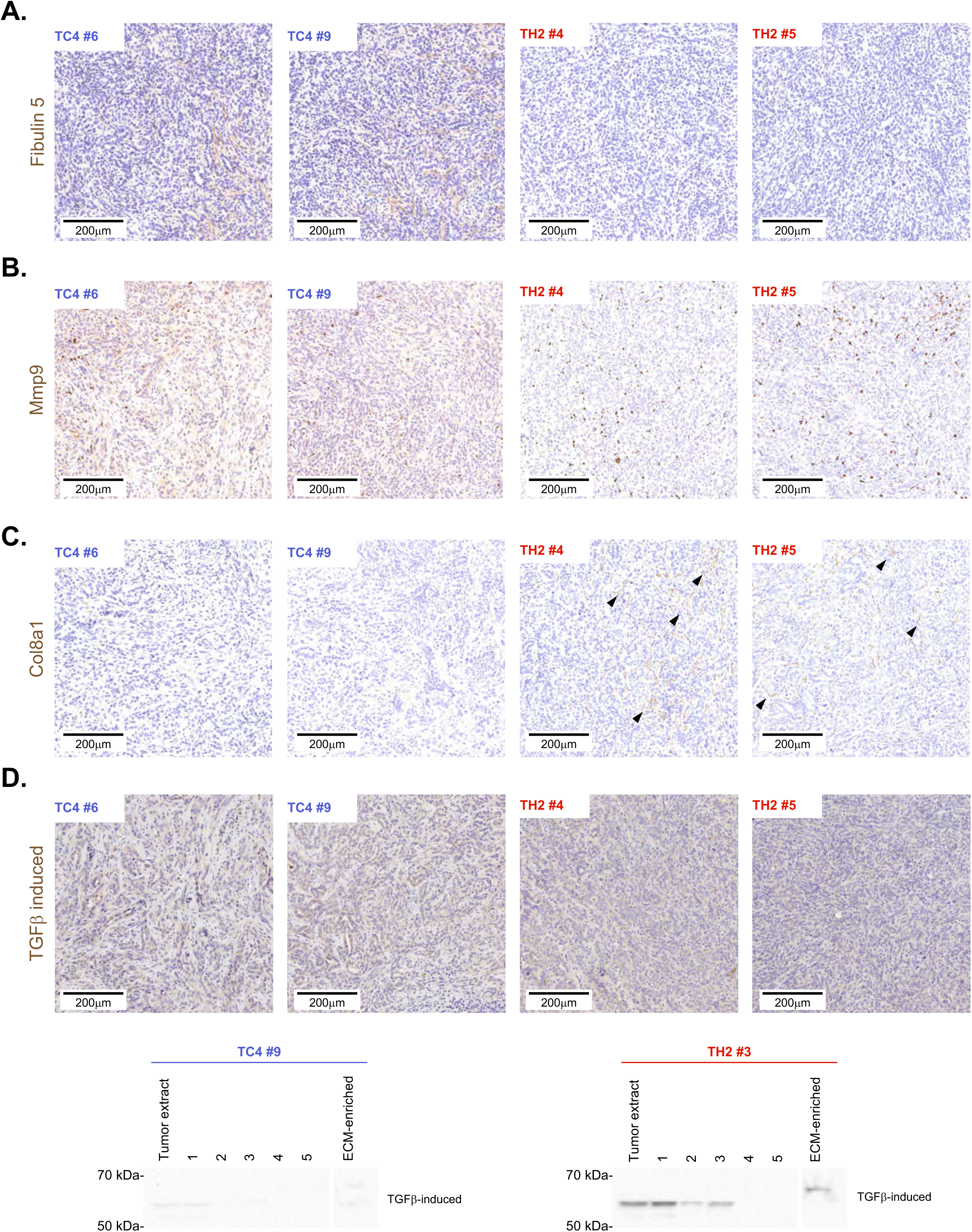
Orthogonal validation of proteins detected in differential abundance between CD8^hi^ and CD8^lo^ KPCY tumors using immunohistochemistry. **A.** Representative images of 5µm-thick sections from two independent CD8^lo^ TC4 and CD8^hi^ TH2 tumors stained with an anti-fibulin 5 antibody, confirming the proteomic data and showing the presence of fibulin 5 in the ECM of CD8^lo^ tumors (*left panels*) and its absence in the ECM of CD8^hi^ tumors (*right panels*) (scale bar = 200μm). **B.** Representative images of 5µm-thick sections from two independent CD8^lo^ TC4 and CD8^hi^ TH2 tumors stained with an anti-Mmp9 antibody confirming the proteomic data and showing the presence of Mmp9 in the ECM of CD8^lo^ tumors (*left panels*) and its absence in the ECM of CD8^hi^ tumors (*right panels; note that the weak positive staining in TH2 tumor sections is intracellular*) (scale bar = 200μm). **C.** Representative images of 5µm-thick sections from two independent CD8^lo^ TC4 and CD8^hi^ TH2 tumors stained with an anti-collagen VIII (Col8a1) antibody confirming the proteomic data and showing the absence of Col8a1 in the ECM of CD8^lo^ tumors (*left panels*) and its presence in CD8^lo^ tumors (*arrow heads, right panels*) (scale bar = 200μm). **D.** *Top panel:* Representative images of 5µm-thick sections from two independent CD8^lo^ TC4 and CD8^hi^ TH2 tumors stained with an anti-TGFβ induced antibody showing positive signals in both CD8^lo^ and CD8^hi^ tumors (scale bar = 200µm). *Bottom panel*: Immunoblots illustrate the abundance of TGFβ induced in protein fractions of increasing insolubility and show an enrichment of TGFβ induced in the ECM-enriched protein fractions of CD8^hi^ TH2 tumors (*right panel*) but not of CD8^lo^ TC4 tumors (*left panel*).

### Signature analysis reveals different pathways activated in CD8^lo^ and CD8^hi^ PDAC tumors

Protein-protein interactions (PPIs) between matrisome components intrinsically determine their functions. Indeed, PPIs allow the assembly of the complexes forming the fibers of the ECM scaffold, while interactions between ECM-remodeling enzymes and their substrates are critical for ECM remodeling. We thus thought to determine whether the components of the matrisome signatures identified in this study participated in similar protein networks using STRING analysis (48). Distinct nodes were identified for the CD8^lo^ and CD8^hi^ matrisome signatures (**Figure 4A, left panel and Figure 4B, left panel, respectively**). However, it is important to note the complete absence of connection between many components, while they all belong to the same compartment. This demonstrates the critical gap in knowledge that remains around the ECM, and highlights the importance of descriptive studies establishing phenotypic correlations that can then enrich the list of signatures in pathway databases (57).

**Figure 4.**
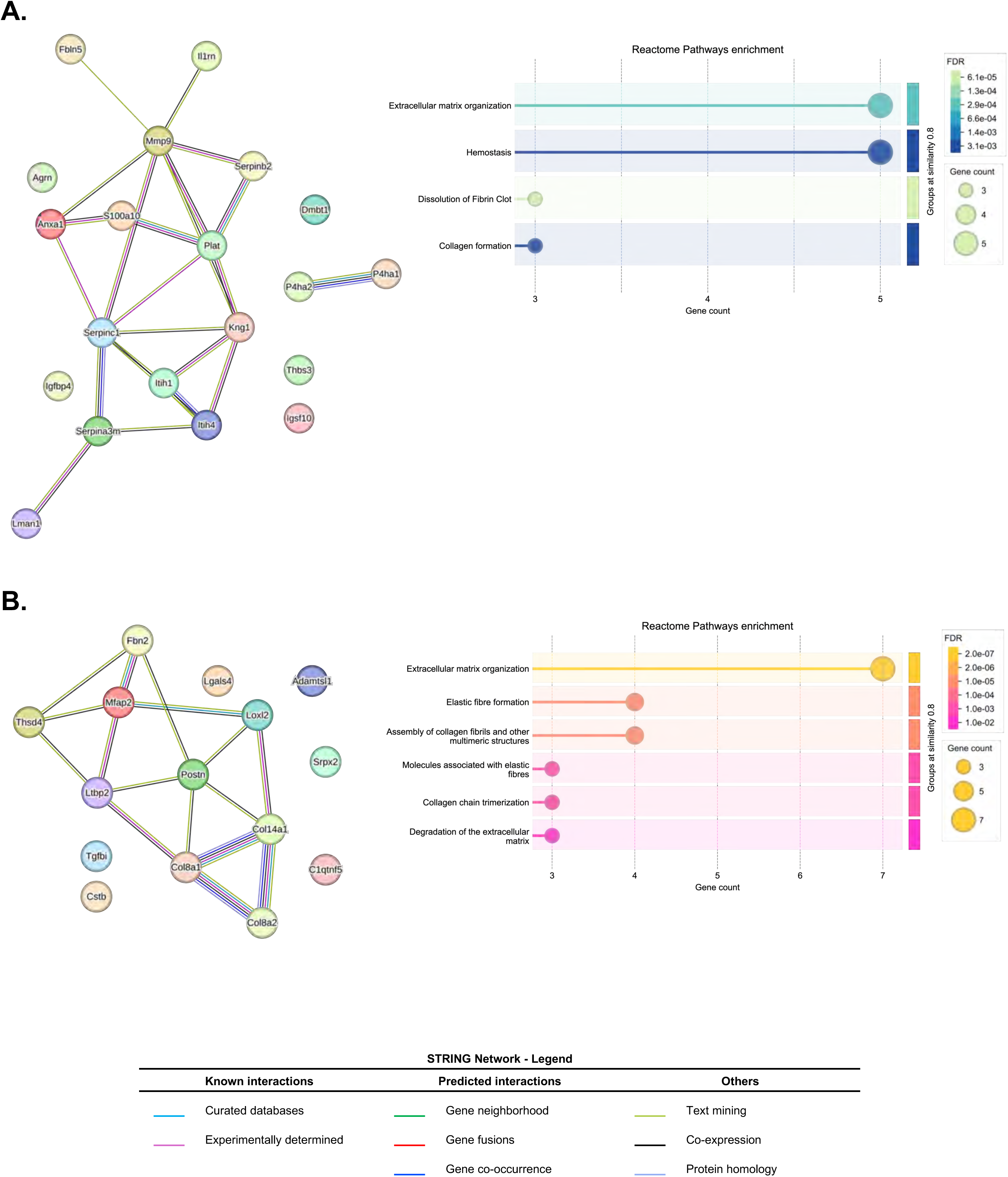
Signature analysis reveals different matrisome networks and pathways activated in CD8^lo^ and CD8^hi^ KPCY tumors. **A.** STRING analysis was applied to build a network connecting the proteins of the CD8^lo^ signature (left panel) and to identify Reactome pathways up-regulated in CD8^lo^ tumors (right panel). **B.** STRING analysis was applied to build a network connecting the proteins of the CD8^hi^ signature (left panel) and to identify Reactome pathways up-regulated in CD8^hi^ tumors (right panel).

Next, we thought to determine whether the components of the matrisome signatures identified participated in common signaling pathways. As expected, and due to our experimental design, both signatures were enriched for genes involved in “extracellular matrix organization” as defined by the Reactome database (58). In addition, the CD8^lo^ (*i.e.*, cold tumor) matrisome signature was enriched for genes involved in pathways modulating collagen formation, in line with the phenotype reported in **Figure 1D**, hemostasis, and dissolution of fibrin clot (**Figure 4A, right panel**), suggestive of a higher vascularization or a leaky vasculature in these tumors. In contrast, the CD8^hi^ (*i.e.*, hot tumor) matrisome signature was enriched for genes playing roles in pathways modulating elastic fiber formation or degradation of the ECM (**Figure 4B, right panel**). Further examination of the ECM signature of CD8^hi^ tumors revealed that several of its components are part of the TGFβ signaling pathway, such as the latent TGFβ binding protein 2, TGFβ induced, and fibrillin 2. It is tempting to speculate that a microenvironment enriched in elastic components is more permissive to immune cell infiltration than a microenvironment rich in fibrillar collagen stabilized by extensive post-translational modifications, as suggested by the ECM protein signature of cold tumors.

### Multiple cell populations of the TME contribute to the human PDAC matrisome

ECM proteins are synthesized intracellularly and secreted into the extracellular space by multiple cell types. The assembled ECM of the tumor microenvironment is thus the product of multiple cell types. While fibroblasts and cells of mesenchymal origins, such as stellate cells in the pancreas, are the main producers of core-matrisome components, many other cell types can contribute to the production of the tumor ECM. We thus sought to determine which cell populations within human PDAC express genes of the cold and hot matrisome signatures. To do so, we leveraged a previously published single-cell RNA sequencing dataset (43) reporting gene expression levels across ten cell populations found in PDAC **(Figure 5A**). As expected, we found that genes encoding the proteins of the CD8^lo^ (cold) and CD8^hi^ (hot) matrisome signatures were expressed by different cell populations within the human PDAC microenvironment. For example, for the genes encoding proteins associated with the low CD8^+^ T-cell infiltration, AGRN is expressed by both ductal cell types 1 and 2 and endothelial cells, FBLN5 is expressed by fibroblasts and endothelial cells, MMP9 is expressed by macrophages, fibroblasts, and ductal cell type 2, THBS3 is expressed by stellate cells and fibroblasts, and S100A10 ductal cell types 1 and 2 and a subset of macrophages (**Figure 5B**). Similarly, for the genes encoding proteins associated with a higher CD8^+^ T-cell infiltration, COL8A1, COL8A2, and LTBP2 are primarily expressed by CAFs, POSTN and TGFBI are expressed by CAFs and a subset of endothelial cells, and TGFBI is additionally expressed by stellate cells (**Figure 5C**), as was LOXL2 (not shown). Although gene expression does not automatically equal protein production, these results demonstrate that multiple cell populations within the human PDAC microenvironment likely contribute to the production of the components of the tumor ECM.

**Figure 5.**
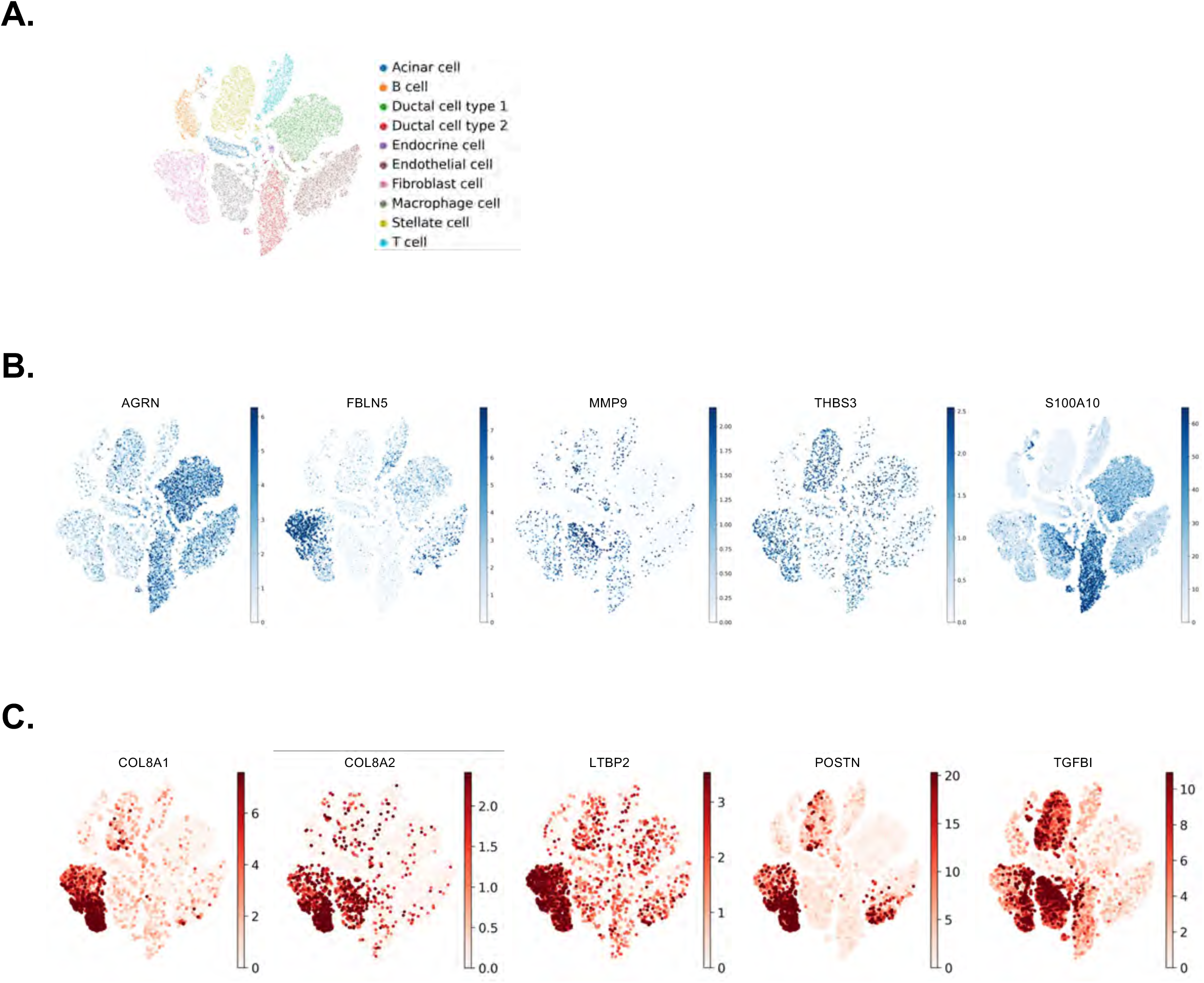
Identification of the cell types producing matrisome proteins in human PDAC microenvironments. **A.** t-SNE represents the distribution of the different cell populations in human PDAC samples. **B.** t-SNE of five genes AGRN, FBLN5, MMP9, THBS3, and S100A10, encoding proteins detected in higher abundance in CD8^lo^ tumors, demonstrates the contribution of different cell populations to the matrisome of these tumors. **C. t-SNE** of five genes, COL8A1, COL8A2, LTBP2, POSTN, and TGFBI, encoding proteins detected in higher abundance in CD8^hi^ tumors, demonstrates the contribution of different cell populations to the matrisome of these tumors.

### Matrisome gene signatures correlate with CD8^+^ T cell infiltration in human PDAC

To further explore the biological relevance of these ECM protein signatures, we examined their correlation with cytotoxic T lymphocyte (CTL) activity. We first performed single-sample gene set enrichment analysis (ssGSEA) to determine the degree to which the hot and cold matrisome gene sets correlated with a gene set indicative of cytotoxic lymphocyte (CTL) infiltration. We found that the hot matrisome gene set showed a moderate positive correlation with the CTL signature (r = 0.33). In contrast, the cold matrisome gene set exhibited a weaker correlation with the CTL signature (r = 0.09) (**Figure 6A**). We next applied the Tumor IMmune Estimation Resource (TIMER) algorithm (45) to determine whether the expression of the genes encoding the ECM protein signatures defined in this study correlated with the abundance of different types of immune infiltrates in human PDAC patient samples of The Cancer Genome Atlas (TCGA PAAD) cohort. We found that the expression of a subset of the genes of the CD8^hi^, hot, PDAC matrisome signature, including COL14A1, ADAMTSL1, LTBP2, FBLN2, and THSD4, positively correlated with the presence of CD8^+^ T cell infiltrates in TCGA PDAC samples, inferred by the TIMER algorithm using defined subsets of biomarkers to annotate CD8^+^ T cells (*e.g.*, TIMER, CIBERSORT, EPIC; **Figure 6B, Supplementary Table S3A**). Conversely, we found that the expression of a subset of the CD8^lo^, cold, PDAC matrisome signature, including S100A10, AGRN, P4HA1, P4HA2, and IL1RN, negatively correlated with CD8^+^ T cell infiltrates (**Supplementary Table S3B**). The fact that there was not complete agreement between the expression of the genes of the gene sets derived from the proteomic signatures defined in this study and the level of immune cell infiltration can partly be attributed to the fact that there is a well-documented discrepancy between RNA expression level and protein abundance. It is also important to note that the data from TCGA include gene expression levels from all the cell types present in tumors, including tumor cells and stromal cells. It would be interesting in the future to correlate matrisome protein abundance in human PDAC samples and the level of immune cell content to further validate the relevance of the protein signature we identified in pre-clinical models of hot and cold PDACs.

**Figure 6.**
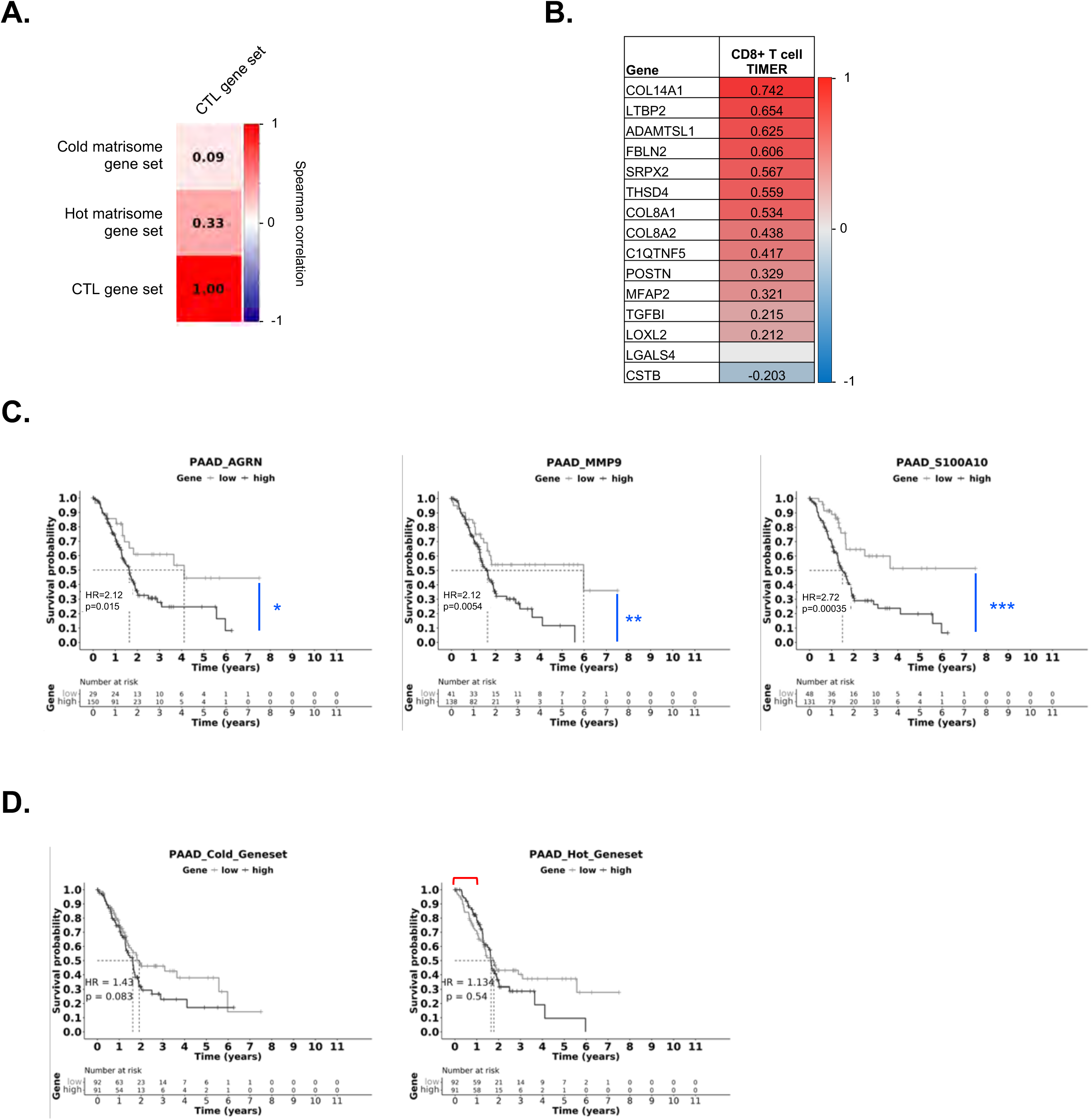
Prognostic value of the expression of matrisome genes on CD8^+^ T-cell infiltration of human PDACs and PDAC patient survival. **A.** Pairwise Spearman correlation coefficients were calculated between the cold ECM gene set or the hot matrisome gene set and a cytotoxic T lymphocyte gene set. Color scale reflects the strength and direction of correlation, with red indicating a positive correlation and blue indicating a negative correlation. **B.** Heat map represents the association between the expression of the genes from the hot PDAC matrisome signatures with CD8^+^ T infiltration assessed using the TIMER2.0 algorithm in the TCGA PDAC RNA-Seq dataset. A positive association score is indicated in red, whereas a negative association is indicated in blue. Associations with a p-value less than 0.05 were considered statistically significant. **C.** Kaplan-Meier plots represent the survival probability (y-axis) over time (x-axis) of PDAC patients stratified with high (black line) or low (grey line) expression levels of the following matrisome genes encoding matrisome proteins characteristic of CD8^lo^ tumors: AGRN, MMP9, S100A10. Statistical analysis was performed using the Cox proportional hazard model, and corresponding hazard ratio (HR) and significance (p-value) are indicated. *: p<0.05, **: p<0.01, ****: p<0.001. **D.** Kaplan-Meier plots represent the overall survival (OS) probability of PDAC patients stratified into two groups based on high expression (black line) or low expression (grey line) scores of the CD8^lo^ matrisome signature (*left panel*) or CD8^hi^ matrisome signature (*right panel*) tumors. Statistical analysis was performed using the Cox proportional hazard model, and corresponding hazard ratio (HR) and significance (p-value) are indicated. Patients expressing a higher level of genes of the entire “hot matrisome signature” had a slightly better overall survival in the first 1.5 years since diagnosis (red bar).

### Matrisome gene signatures correlate with patient outcome

Multiple studies across cancer types have reported a positive correlation between CD8^+^ T-cell infiltration and survival (59–61) or recurrence (62). We thus asked whether the level of expression of genes encoding the proteins of the cold tumor signature negatively correlated with patient survival. To do so, we retrieved expression data and clinical data from The Cancer Genome Atlas and plotted the survival probability of high vs low expressors over time. We found that the level of expression of three of the genes of the cold matrisome signature, AGRN, MMP9, and S100A10, independently correlated with worse prognosis and overall survival (**Figure 6C**). Of note, the expression of the other genes did not correlate with survival (not shown). We also observed that patients expressing a higher level of genes in the entire “cold matrisome signature” had statistically significant worsen overall survival (**Figure 6D, left panel**), while patients expressing a higher level of genes in the entire “hot matrisome signature” had a slightly better overall survival in the first 1.5 years since diagnosis (**Figure 6D, right panel, red bracket**).

## DISCUSSION

Here, we used an ECM-focused proteomic pipeline to identify protein signatures correlating with lymphocyte infiltration in mouse models of PDACs and found that CD8^lo^ – or “cold” – tumors and CD8^hi^ – or “hot” – tumors exhibited different ECM compositions. We further showed that different populations within the tumor microenvironment express the genes encoding these proteins.

SMA^+^ CAFs are commonly thought to be the primary contributors to the production of a collagen-rich ECM within tumors (24,63). Histological analyses show that CD8^lo^ tumors have higher fibrillar collagen content but lower SMA^+^ cell content. Few mechanisms can be proposed to explain this discrepancy: it is possible that CAFs associated with CD8^lo^ tumors produce more fibrillar collagens; other cell populations within the CD8^lo^ TME can also contribute to the production of fibrillar collagens. Last, the collagen ECM in the CD8^lo^ TME could be more stable.

Our proteomic data, showing a higher abundance of enzymes (P4ha1, P4ha2), involved in the hydroxylation of prolines of collagens and the stabilization of triple-helical collagens in the biosynthetic pathway (52), provides support for the third hypothesis. Interestingly, Peng and colleagues have shown that, in melanoma, collagen promotes resistance to immune checkpoint inhibitors (ICIs) and CD8^+^ T cell exhaustion. They further show that decreasing collagen stiffness by inhibiting the collagen crosslinking enzyme lysyl oxidase-like 2 (LOXL2) resulted in increased CD8^+^ T cell infiltration and tumors gained sensitivity to immune checkpoint inhibitors ICIs (64). However, in our dataset, Loxl2 was detected in greater abundance in hot tumors, suggesting that different aspects of collagen remodeling, stabilization via hydroxylation *vs*. crosslinking, may be at play in different cancer types. Nonetheless, it is tempting to speculate that modulating the collagen content in the PDAC TME might alter a tumor’s phenotype and favor immune cell infiltration and activity.

Due to their *extra*-cellular localization and presence in high abundance, ECM proteins are appealing candidates to serve as anchors for the targeted delivery of therapeutic payloads to specific sites such as primary tumors. Indeed, approaches designed to use collagen or tenascin C to anchor and concentrate cytokines or immune checkpoint inhibitors to the tumor ECM have shown promise in animal models (65–69). We thus propose that the proteins of the matrisome signature characteristic of CD8^hi^ tumors defined here could serve as specific anchors to deliver to tumors immune checkpoint inhibitors, since these tumors should readily be able to respond to this modality.

Our findings also underscore the functional relevance of matrisome composition in shaping the immune landscape of tumors. The positive correlation between the hot PDAC matrisome signature, along with enriched expression of markers of CD8⁺ T cells indicative of CTL activity, suggests that these ECM components may actively support T cell infiltration, survival, and effector function. In contrast, the cold matrisome signature is largely absent in T-cell-rich tumor environments, pointing to a role in immune exclusion or suppression. This dichotomy highlights the potential of matrisome profiles not only as biomarkers of immune contexture but also as modulators of antitumor immunity.

The emerging concept of “matritherapy” proposes the development of strategies to modulate either the structure, composition, or signaling functions of the ECM to achieve therapeutic benefits (31). For example, in a recent study, Wu and colleagues showed that knocking down the expression of the ECM protein fibronectin or preventing the interaction of fibronectin with its receptors, α5β1 and αvβ3 integrins, significantly decreased the growth of subcutaneous PDAC in mice (70). Targeting specific matrisome components may thus offer novel therapeutic strategies to reprogram the tumor microenvironment and enhance immune responsiveness, particularly in tumors characterized by a cold, immune-desert phenotype.

## ACKNOWLEDGMENT

The authors would like to thank Dr. Hui Chen and Lasanthi Jayathilaka from the Mass Spectrometry Core facility at UIC and Dr. Maria Sverdlov and Dr. Ryan Easton from the Histology and Tissue Imaging Core facility at UIC for their technical assistance.

## CRediT AUTHOR STATEMENT

**JC:** Investigation, Methodology, Formal Analysis, Visualization, Writing

**DP:** Formal Analysis, Visualization, Writing

**SAO:** Investigation, Resources, Supervision

**JP:** Investigation

**JP:** Conceptualization, Funding Acquisition, Investigation, Resources

**DF:** Investigation, Visualization, Writing

**ASK:** Funding Acquisition, Supervision, Conceptualization, Resources, Project Administration

**NS:** Supervision, Resources, Project Administration, Writing

**AN:** Conceptualization, Methodology, Resources, Formal Analysis, Visualization, Writing, Supervision, Funding Acquisition, Project Administration

## DATA AVAILABILITY STATEMENT

Raw mass spectrometry data and accompanying metadata file in the Sample and Data Relationship Format (SDRF) (49) have been deposited to the ProteomeXchange Consortium (50) via the PRIDE partner repository (51) with the dataset identifier PXD060932. The raw data will be made publicly available upon acceptance of the manuscript.

## SUPPLEMENTARY FIGURE LEGENDS

**Supplementary Figure S1.**
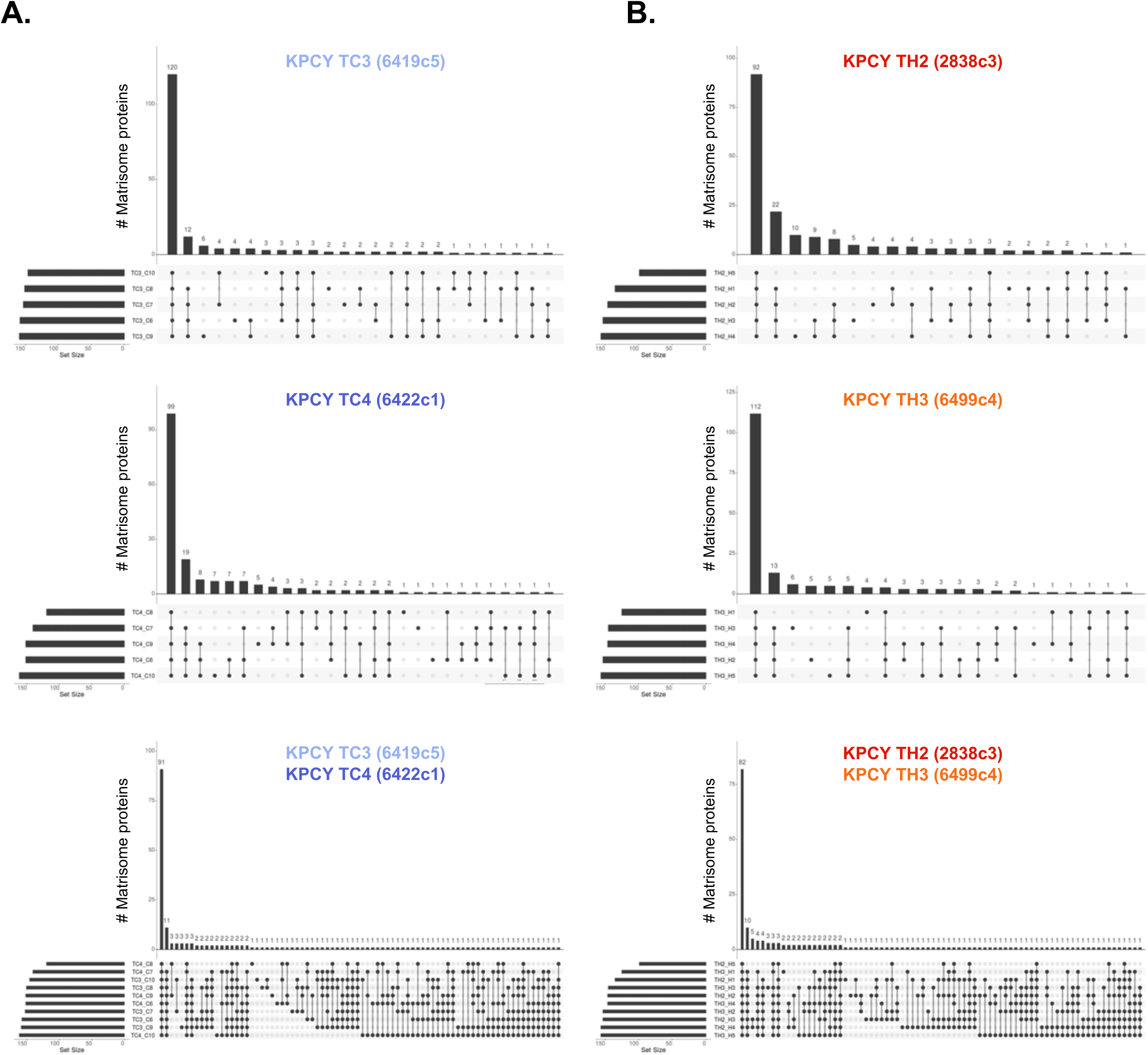
**A.** Upset plots represent the overlap between the matrisome proteins identified by mass spectrometry in CD8^lo^ TC3 (6419c5 cell line; top panel), TC4 (6422c1 cell line; middle panel), or across both TC3 and TC4 KPCY PDAC samples (lower panel). **B.** Upset plots represent the overlap between the matrisome proteins identified by mass spectrometry in CD8^hi^ TH2 (2838c3 cell line; top panel), TH3 (6499c4 cell line; middle panel), or across both TH2 and TH3 KPCY PDAC samples (lower panel).

## SUPPLEMENTARY TABLE LEGENDS

**Supplementary Table S2.** List of antibodies used in this study.

| **Antibody target** | **Source** | **Catalog #** | **Host** | **Application** | **Concentration or Dilution** |
| --- | --- | --- | --- | --- | --- |
| **Actin** | Hynes lab (MIT) | Clone 14-4 | Rabbit | Western blot | 1:5,000 |
| **Collagen I (Col1a1)** | Sigma | AB765P | Rabbit | Western blot | 0.5 µg/mL |
| **GAPDH** | Sigma | MAB374 | Mouse | Western blot | 2.0 µg/mL |
| **Histone H4** | Sigma | 05-858 | Rabbit | Western blot | 1:30,000 |
| **Integrin β1** | Hynes lab (MIT) | ROH 210 | Rabbit | Western blot | 1:1,000 |
| **αSMA** | Abcam | ab5694 | Rabbit | Immunohistochemistry; HIER | 2.0 µg/mL |
| **Agrin** | Abcam | ab85174 | Rabbit | Immunohistochemistry; HIER Tris, pH 9 | 1.0 µg/mL |
| **CD8** | Cell Signaling | 98941 | Rabbit | Immunohistochemistry; HIER Tris, pH 9 | 3.2 µg/mL |
| **Collagen VIII (Col8a1)** | Sigma | HPA053107 | Rabbit | Immunohistochemistry; HIER Tris, pH 6 | 1.0 µg/mL |
| **Fibulin 5** | Abcam | ab109428 | Rabbit | Immunohistochemistry; HIER Tris, pH 9 | 0.7 µg/mL |
| **MMP9** | Abcam | ab38898 | Rabbit | Immunohistochemistry; HIER Tris pH 9 | 1.0 µg/mL |
| **Periostin** | Abcam | ab14041 | Rabbit | Immunohistochemistry; HIER Tris, pH 9 | 0.5 µg/mL |
| **TGFβ-induced** | Sigma | HPA017019 | Rabbit | Immunohistochemistry; HIER Tris, pH 6 | 0.2 µg/mL |

\* *HIER: Heat-induced epitope retrieval*

